# MtrA is an essential regulator that coordinates antibiotic production and sporulation in *Streptomyces* species

**DOI:** 10.1101/090399

**Authors:** Nicolle F. Som, Daniel Heine, John T. Munnoch, Neil A. Holmes, Felicity Knowles, Govind Chandra, Ryan F. Seipke, Paul A. Hoskisson, Barrie Wilkinson, Matthew I. Hutchings

**Affiliations:** School of Biological Sciences, University of East Anglia, Norwich Research Park, Norwich, United Kingdom. NR4 7TJ; Department of Molecular Microbiology, John Innes Centre, Norwich Research Park, Norwich, United Kingdom. NR4 7TJ; School of Molecular & Cellular Biology, Astbury Centre for Structural Molecular Biology, University of Leeds, Leeds, LS2 9JT, UK; Strathclyde Institute of Pharmacy and Biomedical Sciences, University of Strathclyde, 161, Cathedral Street, Glasgow, G4 0RE, UK

## Abstract

*Streptomyces* bacteria make numerous secondary metabolites, including half of all known antibiotics. Understanding the global regulation of secondary metabolism is important because most *Streptomyces* natural products are not made under laboratory conditions and unlocking ‘cryptic’ biosynthetic gene clusters (BGCs) is a major focus for natural product discovery. Production is coordinated with sporulation but the regulators that coordinate development with antibiotic biosynthesis are largely unknown. Here we characterise a highly conserved actinobacterial response regulator called MtrA in antibiotic-producing *Streptomyces* species. We show that MtrA is an essential global regulator of secondary metabolism that directly activates antibiotic production in in *S. coelicolor* and *S. venezuelae*. MtrA also controls key developmental genes required for DNA replication and cell division and we propose that MtrA is the missing link that coordinates secondary metabolism with development in *Streptomyces* species.

## Introduction

*Streptomyces* secondary metabolites account for two thirds of all known antibiotics and numerous other compounds that are used in human medicine as anticancer, anti-parasitic, antiviral and immunosuppressant drugs. Discovery of these natural products peaked in the 1950s but there has been a resurgence of interest in the 21^st^ century, driven by genome sequencing and the increasing threat of drug resistant infections ^1^. Despite their importance however, we still have a poor understanding of how *Streptomyces* bacteria control the production of their secondary metabolites. This is important because ≥75% of their secondary metabolite biosynthetic gene clusters (BGCs) are not expressed in laboratory culture and activating cryptic BGCs could facilitate the discovery of new antibiotics and other useful compounds ^2,3^.

The major way in which bacteria sense and respond to their environment is through two-component systems and several have been implicated in the regulation of antibiotic production in *Streptomyces* species ^4^. Two component systems typically consist of a bifunctional sensor kinase and a cognate response regulator ^5^. The sensor kinase perceives an extracellular signal and activates its cognate response regulator through a two-step phosphorylation. The phosphorylated regulator (RR∼P) brings about a response to the original signal, usually by modulating target gene expression. In the absence of signal, the bifunctional sensor kinase dephosphorylates its response regulator to keep the response switched off ^6^. The delicate balance of kinase and phosphatase activities is crucial in modulating the activity of the response regulator and its target genes during the bacterial cell cycle ^5^. Cross-talk between systems is rare in wild-type cells but removal of a sensor kinase can result in constitutive activation of its response regulator by a non-cognate sensor kinase or by the cellular pool of acetyl phosphate ^7^. Similarly, altering a sensor kinase to block its phosphatase activity might result in a response regulator that cannot be dephosphorylated and is constitutively active ^5^.

Here we report characterisation of the highly conserved actinobacterial response regulator MtrA in the model organism *Streptomyces venezuelae* ^8^. MtrA was first identified as an essential regulator in *Mycobacterium tuberculosis* (<u>M</u>ycobacterium <u>t</u>uberculosis <u>r</u>egulator <u>A</u>) ^9^. *M. tuberculosis* MtrA (TB-MtrA) regulates the expression of *dnaA* and *dnaN*, which are essential for DNA replication, and sequesters the origin of DNA replication, *oriC*, in dividing cells ^10^. TB-MtrA also regulates the expression of cell division genes and interacts directly with the DnaA protein ^10^. MtrA is activated when the MtrB sensor kinase localises to the site of cell division through interaction with FtsI and DivIVA. These data have led to a model in which oscillations in TB-MtrA∼P levels play a key role in cell cycle progression by repressing DNA replication and activating cell division ^10–12^. In the closely related *Corynebacterium glutamicum*, Cg-MtrA controls genes involved in cell wall remodelling and the osmotic stress response ^13,14^ but deletion of the *mtrAB* genes gives rise to elongated cells which are indicative of a defect in cell division ^15^.

*Streptomyces* bacteria are filamentous saprophytes which grow through the soil as branching substrate mycelium that extends at the hyphal tips. Nutrient starvation triggers the production of reproductive aerial hyphae that accelerate DNA replication, generating up to 200 copies of the chromosome in each aerial hypha, before undergoing cell division to form chains of unigenomic spores. Aerial hyphae production and sporulation are coordinated with the production of bioactive secondary metabolites, including antibiotics. *S. coelicolor* has traditionally been used to study development and antibiotic production because it makes pigmented antibiotics and spores but it only differentiates into aerial hyphae and spores on solid agar. *S. venezuelae* has emerged as an alternative model because it completes a full developmental life cycle in liquid growth medium in ∼20 hours ^16^. Here we report that MtrA is essential for the growth of *S. venezuelae* and directly regulates the expression of genes involved in DNA replication, cell division and secondary metabolism. ChIP-seq throughout the *S. venezuelae* life-cycle shows that Sv-MtrA activity displays a biphasic plasticity such that it is active during vegetative growth and sporulation, but inactive during formation of aerial hyphae. We propose that in *Streptomyces* species, MtrA represses DNA replication and cell division during active growth and following sporulation. We also show that Sv-MtrA binds to sites spanning ∼85% of the BGCs in *S. venezuelae* and directly modulates expression of target genes in at least 75% of these. ChIP-seq in *S. coelicolor* yielded similar results and key developmental genes bound by MtrA in both streptomycetes include *bldM, ftsZ, ssgA, smc, smeA, whil, whiB, whiD* and *wblE*. We propose that MtrA is an essential master regulator that coordinates differentiation and secondary metabolism in *Streptomyces* species.

## Results

### MtrA is essential in *Streptomyces venezuelae*

To investigate the function of the MtrAB two component system in *S. venezuelae* we deleted either *mtrA* or *mtrB* ^17^. *mtrB* was deleted easily but all attempts to delete *mtrA* were unsuccessful until we introduced a second copy of *mtrA in trans* suggesting MtrA is essential in *S. venezuelae*. It follows that MtrA must be active in the absence of MtrB, otherwise deleting *mtrB* would be lethal. We propose that deletion of *mtrB* leads to constitutive phosphorylation of MtrA, either by the cellular pool of acetyl phosphate or another sensor kinase. We observed a similar result with *Streptomyces coelicolor* VanRS, where VanR and its target genes are constitutively active in a Δ*vanS* mutant ^7^. Surprisingly, deletion of *mtrB* has no effect on the growth rate of *S. venezuelae* in liquid medium (Figure S1).

### MtrA activity changes during the life cycle of *S. venezuelae*

Microarray data ^18^ shows that *mtrA* expression levels remain fairly constant during the *S. venezuelae* life cycle and we do not detect any significant change in MtrA protein levels during the life cycle using immunoblotting (Figure S2). Attempts to compare levels of phosphorylated and non-phosphorylated MtrA using Phostag were unsuccessful and so, to examine MtrA activity during the *S. venezuelae* life cycle, we performed ChIP-seq at two-hour time points (8-20 hours) from active growth through to sporulation (accession number GSE84311, Table S1). We expressed MtrA-3xFlag *in trans* under the control of its own promoter and deleted the native *mtrA* gene. Survival of this strain shows that MtrA-3xFlag is functional and there is no significant difference in growth rate between this strain and the wild-type (Figure S1). Analysis of the ChlP-seq data using a *P* value > 0.05 revealed that only one target is enriched at all time points, the promoter region of the *ectABCD* operon, which is the BGC for the secondary metabolites ectoine and 5’ hydroxyectoine (5HE). We do not detect ectoine or 5HE in the wild-type or Δ*mtrB* mutant using LCMS with authentic standards so we predict that MtrA represses *ectABCD* expression throughout the life cycle. Analysis of all the MtrA ChIP-seq datasets using *P* > 0.05 shows that the pattern of binding of MtrA is dynamic throughout the lifecycle with peak DNA binding activity seen at 10 and 20 hours, coinciding with mid-exponential growth, and at 20 hours, in spores (Figure S3). Mapping ChIP-seq reads at targets such as the *mtrA* promoter provides a good illustration of this biphasic activity (Figure 1). Differential RNA sequencing shows that the major *mtrAB-lpqB* transcript is leaderless, with an MtrA binding site immediately upstream of the −35 site, suggesting positive autoregulation. A minor transcript starts at −79bp suggesting two promoters might drive expression of the *mtrAB-lpqB* operon (Figure 1).

**Figure 1.**
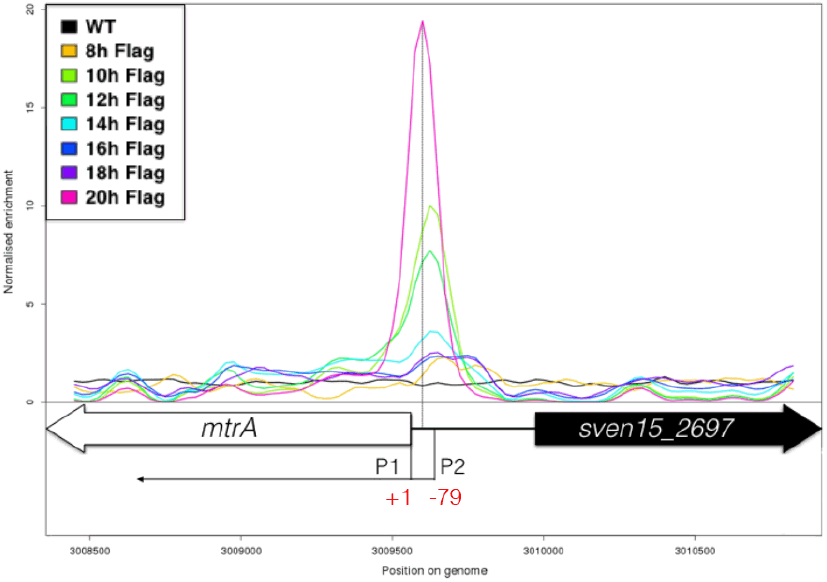
MtrA autoregulates its own expression. ChIP-seq during the S. venezuelae life cycle shows that MtrA has highest activity in actively growing mycelium (10 and 12 hours) and spores (20 hours). The peaks shown are at the mtrA promoter, which has two transcript start sites at +1 (P1) and −79 (P2). Deletion of mtrB increases mtrA transcript levels ∼3-fold indicating positive autoregulation.

#### Identifying the MtrA binding site

To identify an MtrA consensus binding sequence we used MEME ^19^ to analyse three MtrA target promoters identified by ChIP-seq and confirmed *in vitro* using electrophoretic mobility shift assays (EMSAs). This analysis identified an AT-rich 7 bp motif present at all three promoters (Figure 2). We then used MEME to analyse 50 bp of sequence from beneath each peak at the 14h and the 16/18h time points and this analysis identified a conserved sequence at each target which is effectively a direct repeat of the AT rich motif (Figure 2). However, many of the sequences enriched in the MtrA ChIP-seq dataset do not obviously contain this motif and we hypothesise that MtrA may interact with other transcription factors. Immunoprecipitation experiments using anti-Flag beads pulled down four different regulators in the MtrA-3xFlag cultures that are not immunoprecipitated in the wild-type control. The upstream regions of the genes encoding these regulators *(sven15_0243, sven15_2691*, *sven15_3571* and *sven15_4644)* are all bound by MtrA (Table S1). Sven15_3571 is DnaA, which initiates DNA replication and acts as a transcription factor in bacteria, and is also bound *in vivo* by TB-MtrA {Purushotham:2015iq}. Thus, our data suggests (but does not prove) that MtrA forms complexes with other DNA binding proteins and this could explain why there are so many enrichment peaks in the 20-hour ChIP-seq dataset. Interaction between regulatory proteins to form heterodimers is not unprecedented in *S. venezuelae*. The response regulator BldM modulates one set of target genes as a homodimer and another set by forming heterodimers with the WhiI response regulator ^20^. The developmental regulators WhiA and WhiB are also proposed to interact and have identical ChIP-seq regulons ^21^.

**Figure 2.**
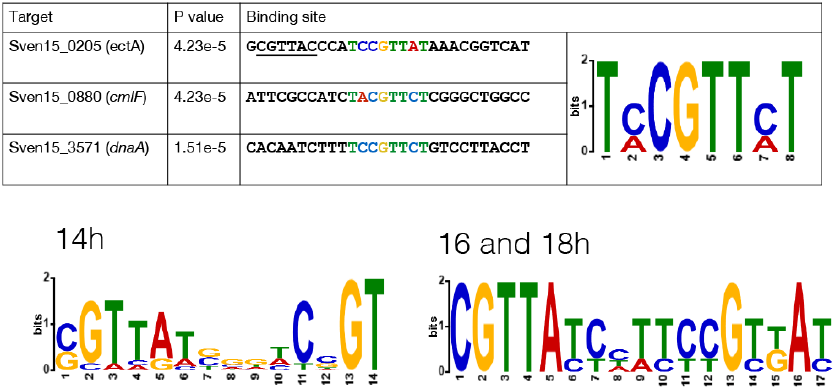
MtrA binds to 6-5-6bp direct repeat motif. The table shows three MtrA ChIP targets that are bound in vitro by purified MtrA protein. They are the promoter regions of ectABCD operon (ectoine BGC), cmlF (chloramphenicol transporter) and dnaA (transcription factor and initiator of DNA replication). MEME analysis identified a conserved 7bp motif (coloured) that is present as a 6-5-6bp repeat motif at the ectA promoter (second half underlined). MEME analysis of different subsets of ChIP targets identified the same direct repeat motif (shown for the targets identified at 14 hours and 16 and 18 hours.

### MtrA regulates the expression of DNA replication and cell division genes

ChIP-seq shows that MtrA activity changes during the *S. venezuelae* life cycle (Figure 1) and RNA sequencing of wild-type and Δ*mtrB* strains shows that expression of *mtrA* is 3-fold upregulated in the Δ*mtrB* mutant (Table S2) which is consistent with activation by MtrA∼P. Given the essentiality of MtrA and its altered activities at different stages of the life cycle we predicted that it could play a key role in regulating cell cycle progression in *S. venezuelae*. Indeed, ChIP-seq shows that many key developmental genes are bound by MtrA in *S. venezuelae* and *S. coelicolor* (see Tables S1 and S2 for complete lists). Remarkably, *M. tuberculosis* and *S. venezuelae* MtrA proteins share some common targets, including the promoter regions of *wblE, dnaA* and *dnaN*, and the *oriC* region between *dnaA* and *dnaN* (Figure 3). RNA-seq shows that deletion of *mtrB* does not affect the expression levels of *dnaA* or *dnaN* under the conditions we used but deletion of *mtrB* causes a two-fold increase in *wblE* transcript levels, suggesting MtrA-dependent activation. MtrA also directly represses *adpA* expression, which is down-regulated nearly 4-fold in the Δ*mtrB* mutant (Table S2).

**Figure 3.**
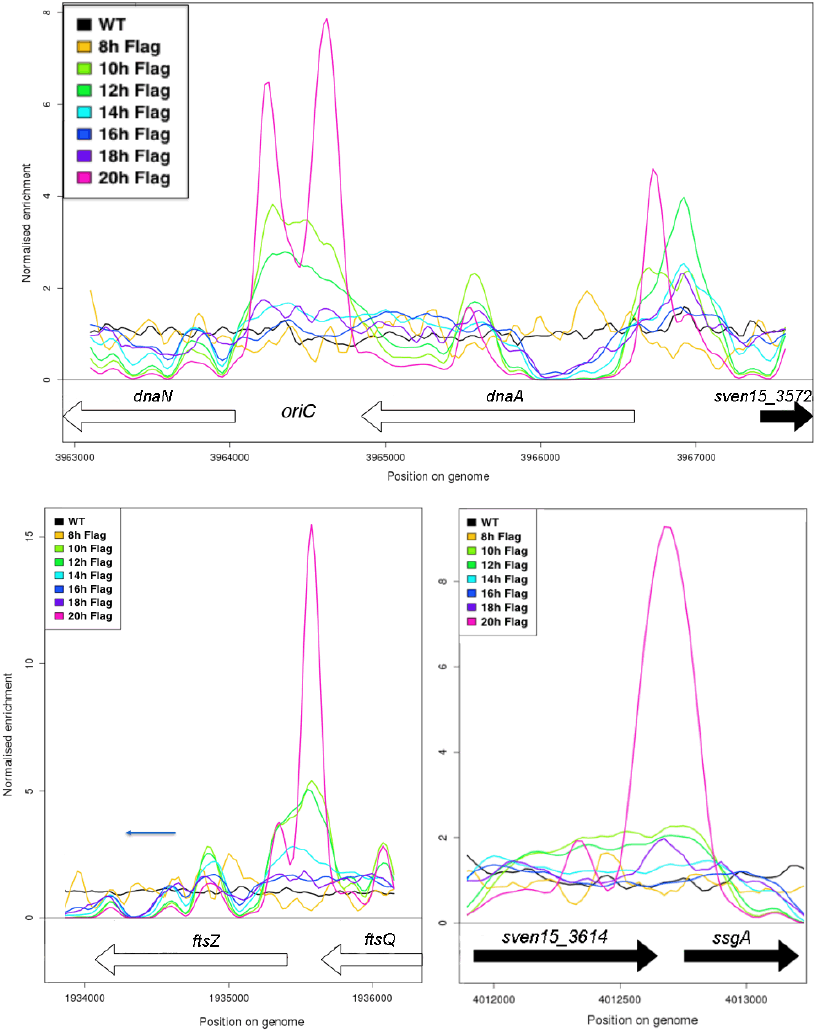
MtrA binds to targets required for DNA replication and cell division in S. venezuelae. Top. MtrA ChIP peaks upstream of dnaA, which encodes the initiator of DNA replication, and dnaN which encodes the DNA polymerase beta clamp subunit and at the origin of DNA replication, oriC. Bottom. MtrA ChIP peaks upstream of ftsZ, which encodes the Z ring forming FtsZ protein and ssgA, whose product helps localise FtsZ to the sites of cell division during sporulation.

AdpA controls key developmental genes and represses DNA replication in *Streptomyces* species ^22,23^. *wblE* encodes a WhiB-like transcription factor of unknown function that appears to be essential ^24^. Sv-MtrA also binds upstream of the cell division genes *ssgA* and *ssgB*, which are upregulated in the Δ*mtrB* mutant, and the *ftsZ* gene, which is unaffected by loss of MtrB, probably because this gene is subject to complex regulation (Figure 3B and Table S2). SsgAB localise the divisome marker protein FtsZ to the correct positions in aerial hyphae to mark the sites of cell division, prior to sporulation ^25,26^. Sv-MtrA also regulates expression of the *smeA-ssfA* operon, which is elevated between 2-and 4-fold in the Δ*mtrB* mutant. SmeA targets the DNA pump SffA to the cell division septa ^16^. MtrA also binds upstream of (and activates some) key developmental regulators, including *whiB, whiD, whiI* and *bldM* (Table S2) ^20,21,27^.

Many of the Sv-MtrA target genes are also repressed by BldD (in complex with c-di-GMP). Sv-MtrA does not regulate *bldD* expression but it does regulate the expression of genes encoding proteins that metabolise cyclic di-GMP ^28^ (Table S2).

### MtrA regulates global BGC expression in *Streptomyces venezuelae*

ChIP-seq shows that MtrA binds to sites spanning 28 of the 31 BGCs predicted by antiSMASH 3.0 in the *S. venezuelae* genome ^29^ (Table S3). The only BGCs with no MtrA enrichment are those encoding biosynthesis of the desferrioxamine siderophores, the WhiE polyketide spore pigment and an unknown natural product. Of these three clusters, the WhiE BGC is upregulated in the Δ*mtrB* mutant suggesting indirect regulation by MtrA, probably via BldM ^20^. The other two BGCs are unaffected by loss of MtrB (Table S3). Of the 28 BGCs that are bound by MtrA, nine have genes that are positively regulated by MtrA, 10 have genes that are negatively regulated by MtrA and three are subject to both positive and negative regulation by MtrA at individual genes within the gene cluster (Table S3). The other six BGCs have sites that are enriched in MtrA ChIP-seq but expression of the genes nearest the ChIP-seq peaks are not affected by deletion of *mtrB* under the conditions we used (Table S3). We predict that one of these, the *ectABCD* operon, is repressed by MtrA because the *ectA* promoter is bound by MtrA at all time points. Consistent with this, we cannot detect ectoine or 5HE in the wild-type or Δ*mtrB* strains. Since most of the *S. venezuelae* BGCs are uncharacterised we know little about their cluster specific regulation, or their products. The antiSMASH predictions we have used here are also likely to include additional flanking genes that may not be part of the BGC so the actual number of BGCs bound by Sv-MtrA may be slightly lower.

### MtrA activates chloramphenicol production in *S. venezuelae*

Phenotypic screening of the Δ*mtrB* mutant revealed that a cryptic antibacterial is activated by removing MtrB but not by simply over-expressing *mtrA* in wild-type *S. venezuelae* (Figure S4).

This is probably because MtrA∼P levels only increase in the absence of MtrB. *S. venezuelae* encodes the biosynthetic pathway for the antibiotic chloramphenicol, which according to previous studies is silent in the wild-type strain ^30^. ChIP-seq shows that Sv-MtrA binds in the intergenic region between the divergent *cmlF* (*sven15_0879*) and *cmlN* (*sven15_0880*) genes (Figure 4) and expression of *cmlN* is 6-fold upregulated in the Δ*mtrB* mutant, consistent with direct activation by MtrA. CmlN is an efflux permease, predicted to export chloramphenicol ^30^. HPLC confirmed that chloramphenicol is produced in the Δ*mtrB* mutant but we also detected very low levels in the wild-type strain suggesting the cluster is not completely silent (Figure 4). Cultivation for 24 hours in biological and technical triplicates confirmed an increased production of chloramphenicol in the Δ*mtrB* mutant with a mean concentration of 0.41 mg/L which is >30 times higher than the wild-type strain (0.013 mg/L) or wild-type over-expressing *mtrA* (0.010 mg/L).

**Figure 4.**
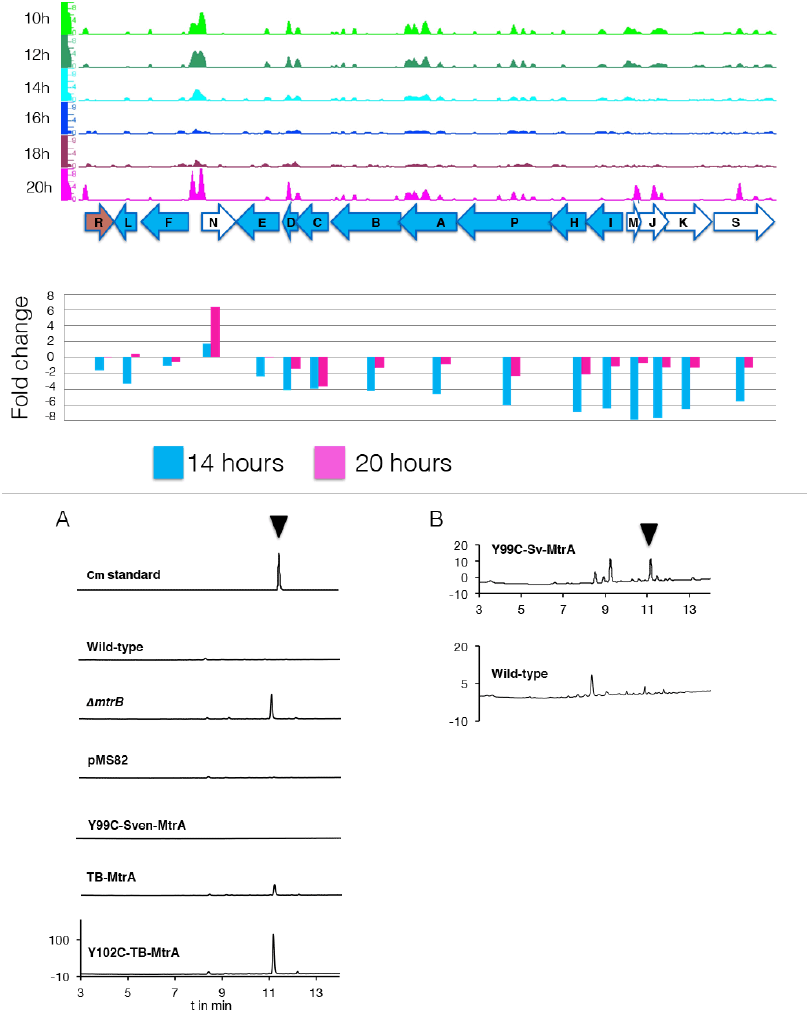
MtrA controls chloramphenicol production. Top. MtrA ChIP peaks at the chloramphenicol (Cm) BGC show two significant peaks between cmlN and cmlF and a smaller peak upstream of the regulator cmlR which is P<0.05. Middle. Expression data for wild-type and ΔmtrB strains at 14 and 20 hours shows cmlF is 6-fold upregulated in ΔmtrB suggesting it is activated by MtrA. All the other Cm genes are downregulated in the mutant which may be due to negative feedback. Bottom left. HPLC shows the Cm standard and extracts of wild-type, ΔmtrB, wild-type plus empty pMS82 vector, and wild-type plus pMS82 expressing Sv-Y99C MtrA, TB-MtrA and TB-Y102C MtrA. Right. Zooming in shows that Sv-Y99C MtrA induces production of low levels of Cm production.

## Deleting *mtrB* has a global effect on the *S. venezuelae* metabolome

While loss of MtrB clearly leads to increased production of chloramphenicol, we were interested to know if there are any other effects on the metabolome. We cultivated the wild type strain and three independently isolated Δ*mtrB* mutants in biological and technical triplicates and analysed the extracts by UPLC/HRMS using an untargeted metabolomics approach. Runs were aligned to compensate for between-run variation and a peak-picking algorithm was applied to allow for the immaculate matching of each feature (a discrete *m/z* value and its retention time) among all runs. Following normalisation, features could be compared quantitatively and their putative identity proposed based on their high-resolution MS-signal. Comparing the level of metabolite signals, it appeared that all Δ*mtrB* mutants showed increased production of a considerable portion of their putative secondary metabolites. To display multidimensional data, we used Principle Component Analysis (PCA) (Figure 5).

**Figure 5.**
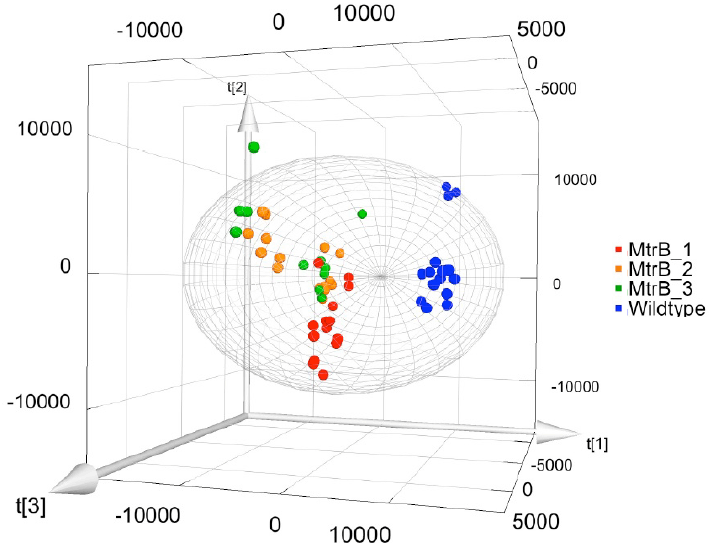
Loss of MtrB results in a global shift in the S. venezuelae metabolome. Principle Component Analysis on the wild-type (blue dots) and triplicate samples of the ΔmtrB strain (red, green and orange dots). Data from ΔmtrB mutant strains clearly group together, and are distinct from data obtained from the wildtype while variations within each group are comparably small.

Each sphere in the 3D Plot represents one dataset obtained from a particular UPLC-HRMS run. Data from the Δ*mtrB* mutant strains clearly group together, and are distinct from data obtained from the wild type, while variations within each group are comparably small. The 3D Plot therefore shows consistent and global changes in the metabolome upon loss of MtrB (Figure 5).

## MtrA directly activates antibiotic production in *Streptomyces coelicolor*

MtrA is conserved in all sequenced *Streptomyces* genomes so we reasoned that it might activate antibiotic production in other streptomycetes. To test this, we deleted *mtrB* in the model organism *Streptomyces coelicolor* ^31^. The 16S rDNA phylogenetic tree of the family *Streptomycetaceae* shows that *S. venezuelae* (clade 40) is highly divergent from *S. coelicolor* (clade 112) so we reasoned that if deleting *mtrB* switches on antibiotic production in these ! distantly related species it is probably universal to all streptomycetes ^32^. The *S. coelicolor ΔmtrB* mutant over-produces the red antibiotic undecylprodigiosin and the blue antibiotic actinorhodin and we also detected significant amounts of metacycloprodigiosin, a potent ! anticancer compound encoded by the undecylprodigiosin BGC but not previously reported ! from *S. coelicolor* (Figure 6). Other secondary metabolites are clearly down-regulated in the Δ*mtrB* mutant, including the siderophores desferrioxamine B and E and germicidin A. We performed ChIP-seq in *S. coelicolor* because its BGCs and their natural products are the most well defined of any *Streptomyces* species ^31,33,34^ The results show that MtrA binds to sites spanning 21 of its 29 BGCs (Tables S4 and S5). MtrA binds upstream of the genes encoding the actinorhodin activator ActII-4 and the undecylprodigiosin activator RedZ (Figure S11). MtrA does not bind to the desferrioxamine BGC *(desABCD)* in *S. coelicolor* (Table S4) but does bind upstream of *sco4394*, which encodes DesR, an iron dependent repressor of desferrioxamine biosynthesis (Figure S5). A single type III polyketide gene, *SCO7221*, is responsible for germicidin biosynthesis ^35^ and it is not regulated by MtrA, so the effect on germicidin biosynthesis must be indirect.

**Figure 6.**
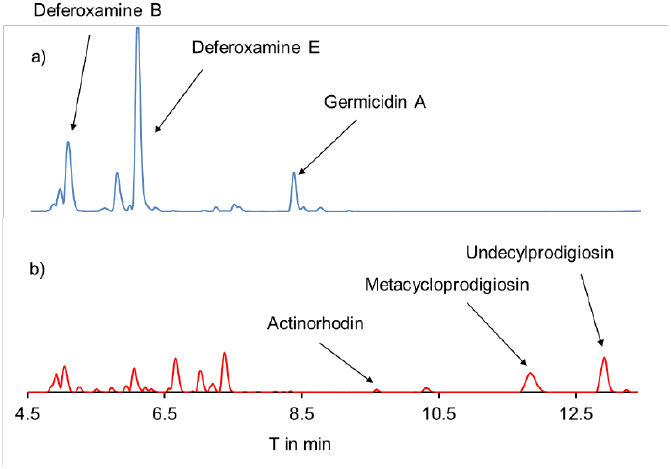
Deleting mtrB switches on antibiotic production in S. coelicolor. (a) HPLC on extracts of wild-type S. coelicolor M145 cultures shows production of desferrioxamines and germicidin. (b) HPLC on extracts of an S. coelicolor ΔmtrB mutant shows production of actinorhodin and undecylprodigiosin is switched on while the production of the desferrioxamine siderophores and germicidin are switched off.

## TB-MtrA activates chloramphenicol production in *S. venezuelae*

Modulating MtrA activity by deleting *mtrB* is time consuming and impractical when applied to multiple *Streptomyces* strains. Instead we tested whether alternative, gain-of-function MtrA proteins might be used to activate antibiotic production. We constructed expression vectors for wild-type TB-MtrA, gain-of-function Y102C TB-MtrA ^36^ and a Y99C Sv-MtrA variant which has the equivalent change to Y102C TB-MtrA (Figure S6). We were unable to delete the native *mtrA* gene in strains carrying these constructs but all three MtrA constructs induced chloramphenicol production in wild-type *S. venezuelae* (Figure 4). Y99C Sv-MtrA switches on the production of relatively small amounts of chloramphenicol in wild-type *S. venezuelae* but the TB-MtrA proteins switch on much higher levels of production, with the strain expressing Y102C-TB-MtrA making more chloramphenicol than the Δ*mtrB* mutant (Figure 4). These results suggest that expressing different MtrA proteins might be a useful avenue to explore in terms of activating BGCs in *Streptomyces* species and our constructs are available from AddGene (IDs 85988-94).

## Discussion

Our data, and previously published work on MtrA in mycobacteria and corynebacteria, support a model in which removing MtrB results in constitutive phosphorylation and activation of MtrA which then constitutively activates or represses its target genes in the cell. In *S. coelicolor* and *S. venezuelae* these include key developmental genes and there is a significant overlap with the regulon controlled by the master repressor of development, BldD, which is activated for DNA binding by the secondary messenger cyclic-di-GMP ^28^. Exactly where MtrA fits in to the hierarchy of developmental regulation is not yet clear but MtrA has an essential role in *S. venezuelae* and shares some functions with *M. tuberculosis* MtrA ^10^. Here we have focused on the role of MtrA in regulating secondary metabolism and how it might be exploited to activate cryptic BGCs. The effects on antibiotic production appear to be direct in both *S. coelicolor* and *S. venezuelae* because MtrA activates expression of the chloramphenicol transporter CmlN and switches on chloramphenicol production. It also binds to the promoters of the cluster specific activators ActII-4 and RedZ in *S. coelicolor* and activates the products under their control, actinorhodin and undecylprodigiosin. In a developmental time course it is remarkable that MtrA binds to sites spanning 27 of the 31 BGCs in *S. venezuelae* and directly affects the expression of target genes in at least 22 of these BGCs. In vegetatively growing *S. coelicolor* MtrA binds to sites spanning 21 out of 29 BGCs. We propose therefore that MtrA is a master regulator of secondary metabolism in *Streptomyces* species. Consistent with this, unbiased metabolomics analysis of *S. venezuelae* shows that deleting *mtrB* results in a global change in the metabolome. Given that secondary metabolite BGCs are often subject to complex and multilevel regulation, and given the energy costs associated with making these natural products, it is perhaps not surprising that we do not see obvious over-production of other compounds in the Δ*mtrB* strains. We predict that deletion of known BGCs in the Δ*mtrB* mutants and/or depletion of *mtrA* mRNA will have positive effects on the production of other secondary metabolites. In summary, our study has revealed an important role for MtrA in the life cycles of streptomycetes and our data suggest it might play a similar role to CtrA in *Caulobacter crescentus*, acting as a master regulator of the cell cycle alongside other master regulators such as BldD. Rapid progress has been made recently in elucidating the roles of developmental regulators using *S. venezuelae* as a model system ^16^. Our study shows that *S. venezuelae* can also be used as a model to study global effects on secondary metabolism and to elucidate the regulatory cascades that link secondary metabolism and differentiation. Understanding these genetic circuits is essential if we are to unlock the full potential of these bacteria.

## Materials and Methods

### Strains, plasmids and primers

The bacterial strains, plasmids and cosmids and primers used in this study are listed in Tables S6-8. The Sv-and TB-MtrA expression vectors have been deposited with AddGene (ID 85988-94). *S. venezuelae* NRRL B-65442 is deposited in the USDA Agricultural Research Services (ARS) Culture Collection (http://nrrl.ncaur.usda.gov/cgi-bin/usda/prokaryote/report.html?nrrlcodes=B%2d65442). Plasmid stocks were prepared using Qiagen miniprep kits and cosmids were prepared as described previously ^37^. Genes were deleted using the ReDirect PCR targeting method ^17^ and an *S. venezuelae* NRRL B-65442 cosmid library provided by Professor Mark Buttner at the John Innes Centre, Norwich. All of the expression constructs used in this work were made by Genscript by synthesising the *mtrA* alleles with or without 3’ tags and then cloning into the required vectors. Liquid cultures of *E. coli* were routinely grown shaking at 220 rpm in Lennox Broth at 37°C. Liquid cultures of *S. coelicolor* or *S. venezuelae* were grown in Mannitol Yeast Extract Malt Extract (MYM) at 30°C, shaking at 220 rpm. Cultures grown on solid MYM agar were grown at 30°C. Spore stocks of *S. coelicolor* were prepared from cultures grown on Soya Flour + Mannitol (SFM) agar. All media recipes have been published previously and *Streptomyces* spores were prepared as described ^37^. To determine the developmental growth in liquid culture *S. venezuelae* and mutant strains were grown, shaking in 35ml MYM in 250ml conical flasks containing springs at 30°C at 220rpm. A spore inoculum sufficient to reach an OD_600_ of 0.35 after 8 hours of growth was added to 35ml of MYM media in 250 ml flasks containing springs. The culture densities were measured at OD_600_. Development in liquid cultures was monitored using an GXML3000B microscope from GX optical. Pictures of agar plate grown colonies were taken with a Zeiss SVII stereo microscope. SEM images were taken at the bioimaging facility at the John Innes Centre.

### Purification of MtrA-His

The *mtrA* gene was cloned into pETDuet1 (Novagene) to express the protein with a C-terminal hexa-His tag and purified using a batch method. Cell pellets were resuspended in 25ml lysis buffer (75mM Tris-HCl pH8, 20mM NaCl, 0.1% Triton X100, 50 μl 10mg/ml lysozyme, 3 × Pierce EDTA-free Protease Inhibitor Mini Tablets (Thermo Scientific) in 1L) and incubated for 30 minutes at room temperature. The cell lysate was sonicated 2 × 40 seconds at 50Hz with 1 minute in between sonication steps. Cell debris was removed by centrifugation at 18,000rpm for 20 minutes at 4°C in Beckman Coulter Avanti^®^ J-20 high performance centrifuge using a JLA-25-50 rotor (Beckman Coulter). The supernatant was transferred in a fresh 50ml Falcon tube and 350μl of Ni-NTA agarose beads (Qiagen) were added and incubated under gentle agitation for 1 hour at 4°C. The Ni-NTA agarose beads were spun down gently (maximum of 1200rpm, 4°C) and the supernatant was discarded. The Ni-NTA beads were resuspended in 2ml wash buffer (75mM Tris-HCl pH8, 200mM NaCl, 10% glycerol, 10mM MgCl_2_, 0.1mM DTT, 20mM 2-mercaptoethanol in 1L) and transferred in Polypropylene columns (1ml, Qiagen). The beads in the column were washed with 20ml wash buffer. The protein was eluted from the beads with 2.5ml elution buffer (wash buffer plus 350mL imidazole in 1L). Polyclonal antiserum was raised by Cambridge Research Biochemicals. Immunoblotting was performed as described previously ^38^.

### DNA binding studies

Chromatin Immunoprecipitation followed by sequencing (ChIP-seq) was performed on *S. coelicolor* M145 Δ*mtrA* + MtrA-3xFlag and *S. venezuelae* NRRL B-65442 Δ*mtrA* + MtrA-3xFlag grown in liquid MYM medium ^37^. Note that *S. coelicolor* M145 grows vegetatively and does not differentiate under these growth conditions whereas *S. venezuelae* NRRL B-65442 undergoes a full life cycle in 20 hours. Samples were taken at 16 and 20 hours for *S. coelicolor* and at 8, 10, 12, 14, 16, 18 and 20 hours for *S. venezuelae*. ChIP-seq and electrophoretic mobility shift assays (EMSAs) were performed as described previously ^39^ using probes generated by PCR with 6-Fam labelled primers from Integrated DNA Technologies (Table S8). ChIP-seq Gels were visualised using a Typhoon FLA 9500 laser scanner (GE Healthcare) with LBP/BPB1 emission filter, Exmax 495nm Emmax 576nm, at 50μM resolution. ChIP-seq data was analysed as described previously ^40^ and peaks were visually inspected using integrated genome browser ^41^. Binding sites were identified using MEME ^19^.

### RNA-sequencing

Duplicate wild-type and Δ*mtrB* cultures were grown for 14 and 20 hours in MYM medium and RNA was prepared as described previously ^42^. Libraries were prepared and sequenced by Vertis Biotechnologie and analysed using CLC Genomics Workbench.

## LCMS and PCA

Analytical HPLC was performed on an HPLC 1100 system (Agilent Technologies) connected to a Gemini^®^ 3 μm NX-C18 110 Å, 150×4.6 mm column (Phenomenex) and on a Synergi™ 4 μm Fusion-RP 80 Å LC column 250×10 mm. UPLC-MS for metabolic profiling was performed on a Nexera X2 liquid chromatograph (LC-30AD) LCMS system (Shimadzu) connected to an autosampler (SIL-30AC), a Prominence column oven (CTO-20AC) and a Prominence photo diode array detector (SPD-M20A). A Kinetex^®^ 2.6 μm C18 100 Å, 100×2.1 mm column (Phenomenex) was used. The UPLC-System was connected with a LCMS-IT-TOF Liquid Chromatograph mass spectrometer (Shimadzu). UPLC-HRMS Data was acquired on an Acquity UPLC system (Waters Corporation) equipped with an ACQUITY UPLC^®^ BEH 1.7 μm C18, 1.0 × 100 mm column (Waters Corporation) and connected to a Synapt G2-Si high resolution mass spectrometer (Waters Corporation). A gradient between mobile phase A(H_2_O with 0.1 % formic acid) and mobile phase B (acetonitrile with 0.1 % formic acid) at a flow rate of 80 μL/min was used. Initial conditions were 1 % B for 1 min, ramped to 90 % B within 6 minutes, ramped to 100 % B within 0.5 min, held for 0.5 min, returned to 1 % B within 0.1 min and held for 1.9 min. MS spectra were acquired with a scan time of one second in the range of *m/z* = 150–1200 in positive MSe-Resolution mode. The following instrument parameters were used: capillary voltage of 3 kV, sampling Cone 40, source offset: 80, source temperature of 120 °C, desolvation temperature 350 °C, desolvation gas flow 800 L/h. A solution of sodium formate was used for calibration. A solution of leucine encephalin (H_2_O/MeOH/formic acid: 49.95/49.95/0.1) was used as lock mass and was injected every 15 sec. The lock mass has been acquired with a scan time of 0.3 sec and 3 scans were averaged each time. The lock mass (m/z = 556.2766) has been applied during data acquisition.For processing metabolomics data we used the Software Progenesis QI (Waters). All solvents for analytical HPLC and UPLC-MS were obtained commercially at least in HPLC grade from Fisher Scientific and were filtered prior to use. Formic acid (EZInfo 3.0 (MKS Umetrics AB) was used for plotting PCA data.

## Acknowledgements

We thank Kim Findlay for the Scanning Electron Microscopy, Mark Buttner for the gift of *S. venezuelae* NRRL B-65442, cosmid Sv-6-A04, access to microarray data prior to publication and helpful discussions, Matt Bush for critical reading of the manuscript and helpful discussions and Elaine Patrick for technical support. This research was supported by a BBSRC PhD studentship to NS, a UEA-funded PhD studentship to FK, MRC Milstein award G0801721 to MIH, Norwich Research Park translational funding, NPRONET Proof of Concept funding and Natural Environment Research Council responsive mode grants NE/M015033/1 and NE/M014657/1 to MIH and BW and NE/M001415/1 to PAH.

## Author contributions

FK and RFS constructed the *S. coelicolor* mutants and FK undertook phenotype analysis, NS constructed all the *S. venezuelae* strains and constructs, analysed phenotypes and performed ChIP-seq experiments, JTM performed RNA-seq and dRNA-seq experiments, JTM and GC analysed the ChIP-and RNA-seq datasets, NH made constructs and prepared strain extracts for LCMS, DH purified and analysed all natural products including LCMS and PCA analyses, PAH, MIH and BW conceived the study and all the authors analysed data and wrote the manuscript.

